# T-cell Repertoire Characteristics of Asymptomatic and Re-detectable Positive COVID-19 Patients

**DOI:** 10.1101/2021.03.03.433579

**Authors:** Jianhua Xu, Yaling Shi, Yongsi Wang, Yuntao Liu, Dongzi Lin, Jiaqi Zhang, Jing Lin, Wei Hu, Haolan He, Wei Wang, Wentao Fan, Linlin Li, Hai Lan, Chunliang Lei, Kejian Wang, Dawei Wang

## Abstract

**Background:** The prevention of COVID-19 pandemic is highly complicated by the prevalence of asymptomatic and recurrent infection. Many previous immunological studies have focused on symptomatic and convalescent patients, while the immune responses in asymptomatic patients and re-detectable positive cases remain unclear.

**Methods:** Here we comprehensively analyzed the peripheral T-cell receptor (TCR) repertoire of 54 COVID-19 patients in different phases, including asymptomatic, symptomatic, convalescent and re-detectable positive cases.

**Results:** We found progressed immune responses from asymptomatic to symptomatic phase. Furthermore, the TCR profiles of re-detectable positive cases were highly similar to those of asymptomatic patients, which could predict the risk of recurrent infection.

**Conclusion:** Therefore, TCR repertoire surveillance has the potential to strengthen the clinical management and the immunotherapy development for COVID-19.

**Funding:** The Science and Technology Innovation Project of Foshan Municipality (2020001000431) and the National Key Research and Development Project (2020YFA0708001).

## Introduction

As a highly infectious virus, the severe acute respiratory syndrome coronavirus 2 (SARS-CoV-2) caused the pandemic of coronavirus disease 2019 (COVID-19) (1, 2). Despite the implementation of preventive measures, COVID-19 is spreading worldwide and has affected over 20 million people to date (3). The clinical manifestations of infected patients ranged from asymptomatic condition to severe symptoms (4). And among the COVID - 19 convalescent patients, some tested positive again after discharge (i.e., re-detectable positive) (5).

The global efforts to end COVID-19 are complicated by the prevalence of asymptomatic and recurrent infection. Many previous immunological studies have focused on symptomatic and convalescent patients (6–13), while the immune responses in asymptomatic patients and re-detectable positive cases remain unclear (14, 15). Unlike symptomatic patients who can be effectively identified by clinical features, asymptomatic carriers may inadvertently transmit virus to close contacts and reshape the dynamics of infection in population (16, 17). Although most re-detectable positive cases have minor symptoms and hardly disease progression upon re-admission, their potential infectivity and immunological characterization remain undefined (18, 19).

The antiviral adaptive immunity is greatly dependent on the activation of T-cells, which can selectively eliminate virus-infected host cells (20). The specificity toward viral antigens is determined by the structure of T-cell receptor (TCR) repertoire (21). Based on the advances in sequencing technologies, the biased TCR repertoire in various infectious diseases has be revealed (22–24). Thus, in-depth study on the TCR characteristics in COVID-19 is critically needed (25, 26).

To address the above issues, we conducted a detailed analysis on the peripheral blood samples collected from 54 COVID-19 patients in different phases (including asymptomatic, symptomatic, convalescent and re-detectable positive cases) along with 16 healthy donors. By performing high-throughput sequencing, we explored the TCR repertoire associated to the disease trajectories of COVID-19. Our results presented the unique immunological features of asymptomatic patients and re-detectable positive cases, which could provide help for clinical management and therapy development.

## Results

### Demographic and clinical characteristics

A total of 54 patients with laboratory-confirmed COVID-19 disease were enrolled, including 11 asymptomatic (ASY), 19 symptomatic (SYM), 14 convalescent (CON) and 10 re-detectable positive (RDP) cases. And 16 healthy donors (HD) were recruited as the control group. As shown in Table 1, no significant difference in age or sex was identified between HD group and any of the patient groups (as measured by two-tailed Mann–Whitney U-test or Fisher’s exact test). Except for one patient with pre-existing chronic pharyngitis, most ASY cases had no obvious clinical symptoms during the whole disease course. All HD subjects and most ASY patients had no underlying comorbidity, while some patient of SYM, CON and RDP groups were diagnosed with hypertension, hyperlipemia, diabetes mellitus, cardiovascular disease, chronic liver disease or chronic kidney disease (27). The routine laboratory test results (supplementary Table 1) showed that the lymphocyte levels in the SYM and CON groups were significantly lower than that in the HD group (two-tailed Mann–Whitney U-test p < 0.05). However, such difference was not observed in ASY or RDP group.

**Table 1.**
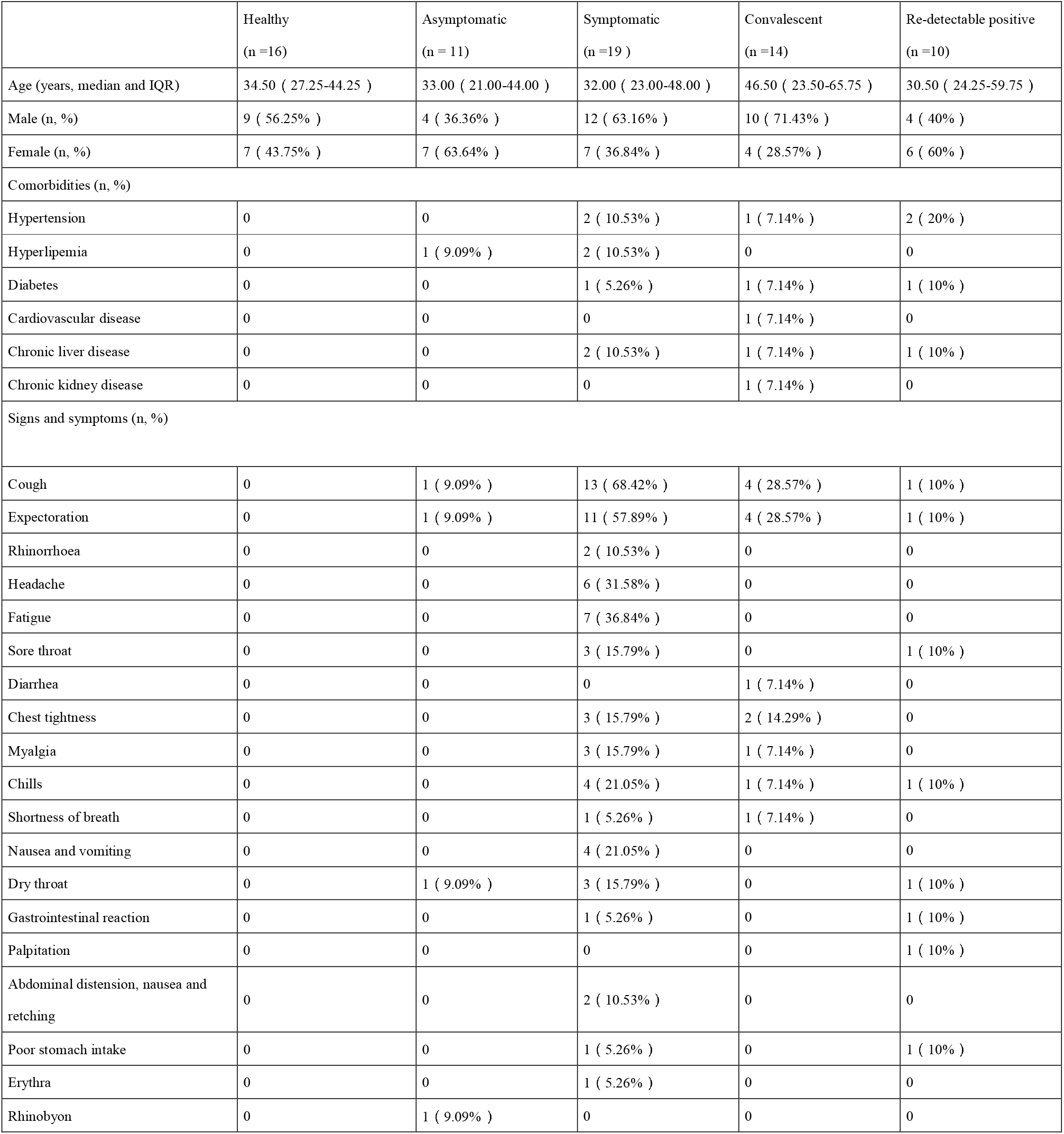
Baseline characteristics of study subjects.

### Patients in different phases have heterogeneous TCR characteristics

Among all the clonotypes identified in our samples, only a small fraction was shared by different groups (Figure 1A). No significant difference in clonal diversity (as measured by D50 index, Shannon entropy and accumulative frequencies of the top 10 clones) was detected between any two groups (Figure 1B-D), suggesting that COVID-19 might not necessarily induce a fundamental change in TCR repertoire composition.

**Figure 1.**
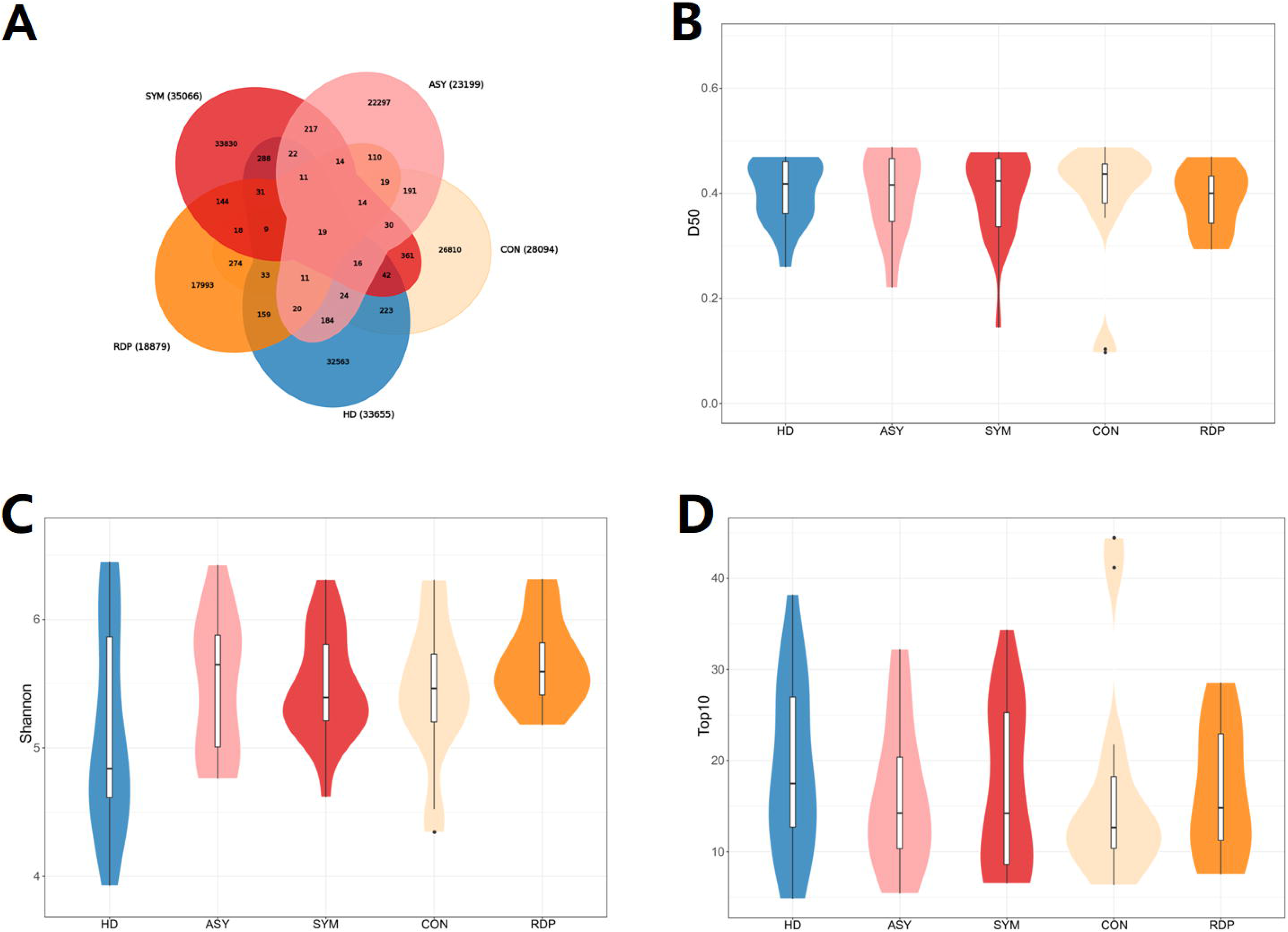
Analysis of overall clonal diversity. (A) The number of overlapped clonotypes among different groups. (B) Distribution of D50 index. (C) Distribution of Shannon entropy. (D) Distribution of accumulative frequencies of the top 10 clones.

In the context of V-J gene usage, we assumed that the dissimilarity between samples can be measured by the Spearman’s correlation coefficient of V-J combination profiles (i.e., stronger positive correlation indicates lower dissimilarity). The dissimilarity between ASY and HD subjects was significantly lower than that between SYM and HD ones (Figure 2A), which demonstrated that asymptomatic infection could be a less aversive stimulus on immune system than symptomatic phase. In addition, RDP showed significantly lower dissimilarity to ASY than to other groups (Figure 2B), suggesting potential commonality between asymptomatic and recurrent infections.

**Figure 2.**
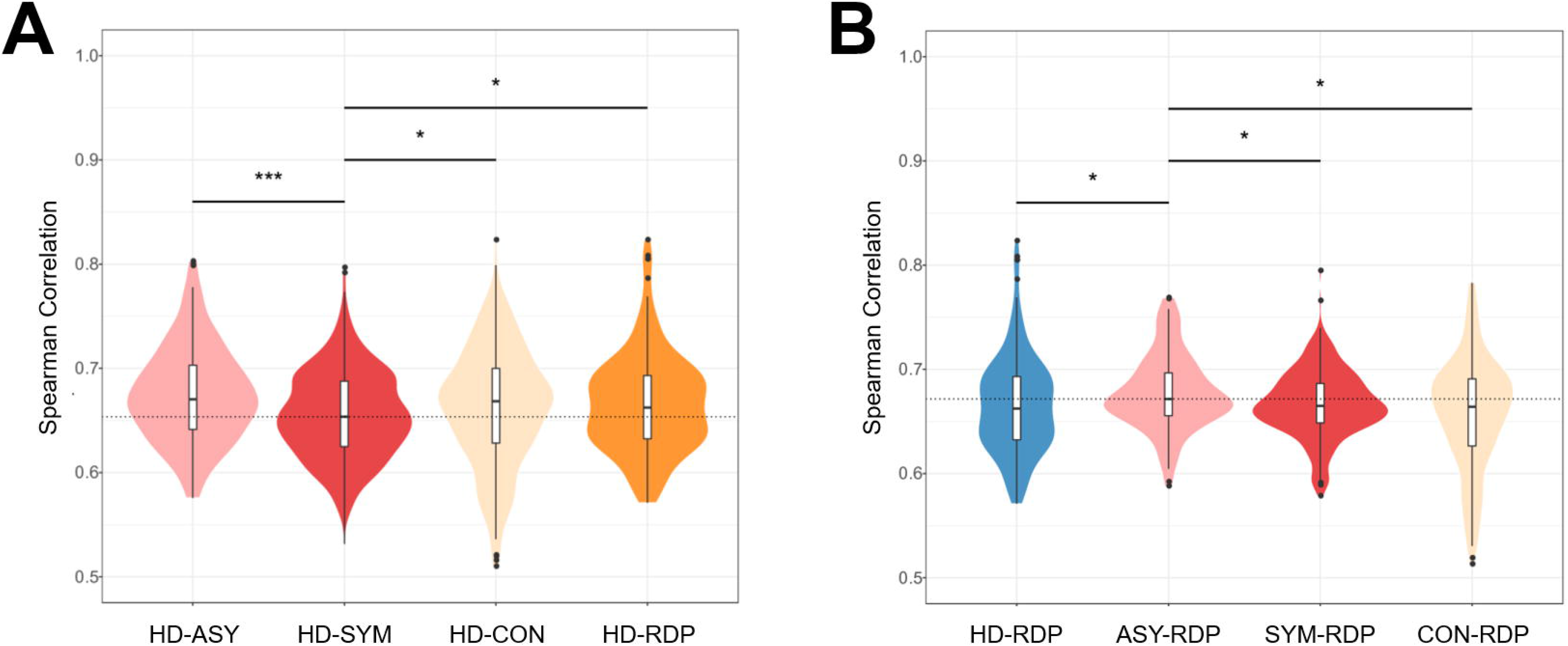
Spearman’s correlation coefficients of V-J combination profiles between different groups. (A) A stronger correlation between HD and ASY groups than that between HD and SYM groups. (B) A strong correlation between RDP and ASY groups.

### Differential gene usage in different phases of COVID-19

Since the immune responses could progress from ASY to SYM group, we further analyzed the differential V-J gene usage. A total of 73 V-J pairings exhibited monotonic trend for increasing or decreasing frequency across HD, ASY and SYM groups (Cuzick’s test for trend p < 0.05, Figure 3A and supplementary Table 2) (28), which illustrated the details of the progressive immune responses. Of note, a significant portion (28.8%, hypergeometric test p = 1.56×10^−12^) of these 73 V-J pairings also showed differential usage between RDP and HD groups (Mann–Whitney U test p < 0.05, Figure 3B and supplementary Table 3), suggesting that certain aspects of adaptive immune responses to SARS-CoV-2 could last till recovery and be awakened upon recurrent infection.

**Figure 3.**
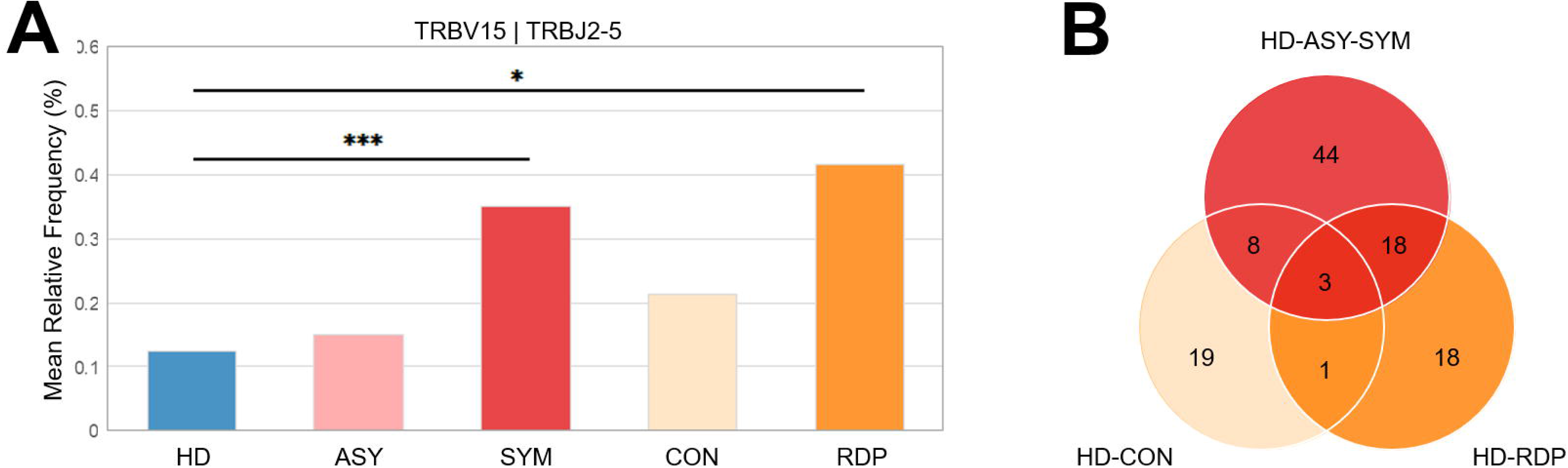
Differential V-J gene usage between different groups. (A) A representative V-J combination showing monotonic increase across HD, ASY and SYM groups. (B) Among the V-J combinations with monotonic changes along the ASY-to-SYM progression, many also showed differential usage in either RDP cases or CON patients as compared to HD subjects.

Since HLA molecules play a crucial role in shaping the TCR repertoire (29), we also sought to determine whether the frequency of certain HLA allele was imbalanced in different groups. By performing Fisher’s exact test following a previously published procedure (22), we confirmed that none of the HLA alleles had significant difference in frequency across groups (Figure 4 and supplementary Table 4), which consolidated our findings in TCR repertoire.

**Figure 4.**
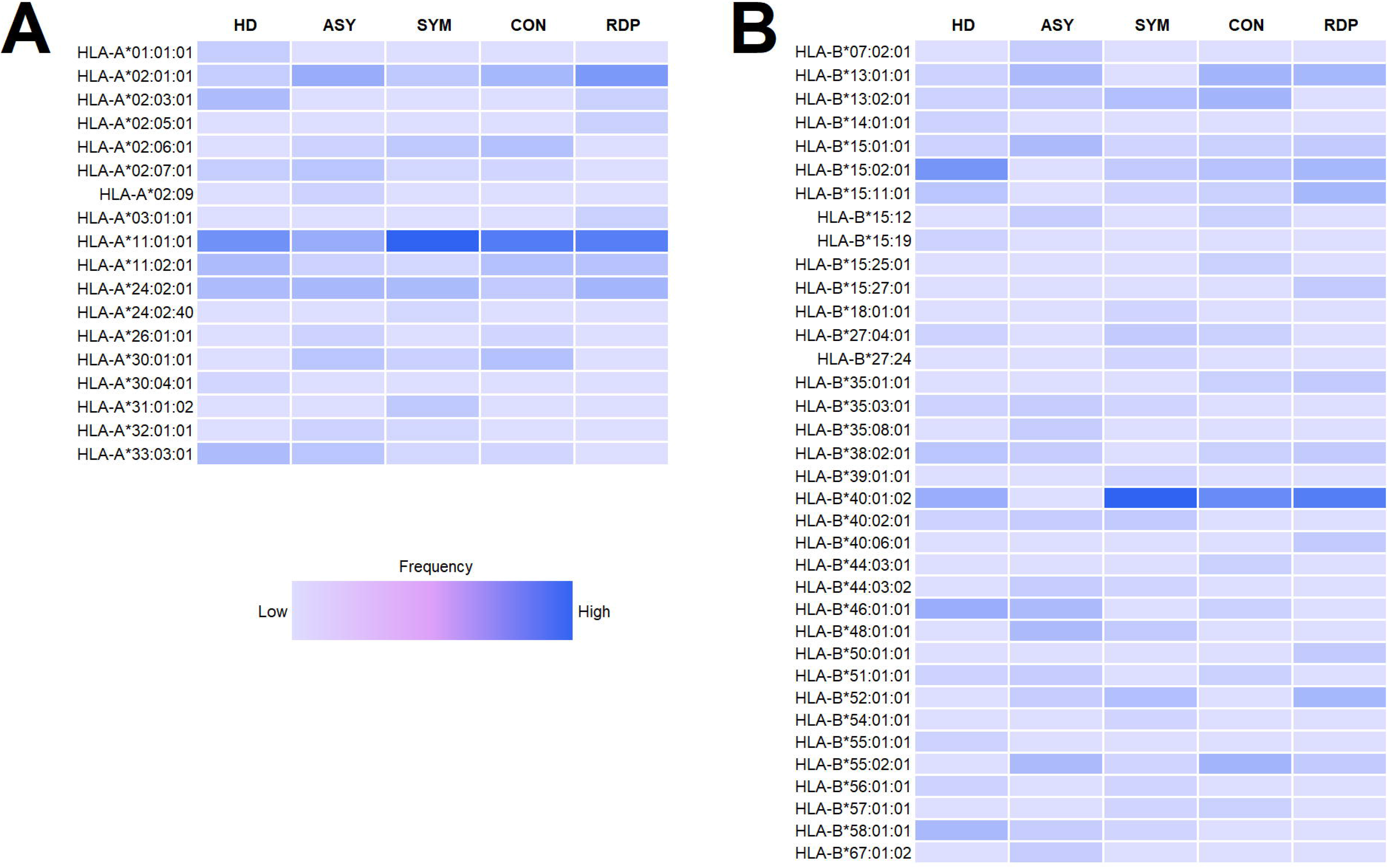
Frequency of HLA-A (A) and HLA-A (B) alleles in different groups.

## Discussion

Asymptomatic and recurrent infections further increased the difficulty of COVID-19 prevention. Here we presented an immunological landscape of COVID-19 patients in different phases, by investigating not only SYM and CON patients but also ASY and RDP ones. Our results demonstrated the divergent trajectories of COVID-19 and particularly the unique characteristics of ASY and RDP patients.

Despite numerous recent studies on COVID-19 patients, our analysis adds understanding of the immune responses to SARS-CoV-2 in several aspects. First of all, it has been repeatedly reported that clinical manifestations tend to vary between asymptomatic and symptomatic patients (14, 30-32). In agreement with such observation, we corroborated that the usage of certain V and J gene segments exhibited monotonic trend across HD, ASY and SYM groups, indicating a possible immune progression from asymptomatic to symptomatic phase. And those V-J combinations showing monotonic changes along the ASY-to-SYM progression can help narrow down the search scope of virus-associated TCR signatures. Identification of TCR specific to SARS-CoV-2 antigens may facilitate the development of highly targeted immunotherapies, such as engineered T-cells for infusion(33, 34)

Next, the risk factors of recurrent infection are not yet well defined (15), which urges development of useful biomarker to enhance clinical management. We found that many V-J combinations with monotonic changes along the ASY-to-SYM progression also showed biased usage in re-detectable positive cases. Such observation indicated the possibility of examining immune repertoire as a measure of clinical outcome and an early predictor of high-risk individuals.

Several limitations of our study should be taken into consideration. Firstly, multiple phases of COVID-19 were characterized by blood samples collected from different individuals, rather than sequential samples from the same infected patients. Therefore, additional longitudinal studies are needed to calibrate the immunological trajectories of COVID-19. Secondly, the enrolled patients were treated with diverse therapies and medications, which may have divergent immunomodulatory effects. Subsequent efforts will optimally require well-controlled clinical trials with unified treatment protocol. In addition, due to limited volume of blood that could be collected in isolation wards, the HLA typing was not performed by DNA sequencing or PCR but estimated with mRNA sequencing data. Thus, the uncertain relevance between HLA alleles and TCR repertoire in COVID-19 should be further explored.

Altogether, this study provided novel insights into the TCR profiles of COVID-19 patients in different phases. A better understanding of the adaptive immunity in COVID-19 could strengthen clinical management and immunotherapy development. Further analyses on large-scale immune repertoire datasets (35, 36) would be required to extend our findings.

## Methods

### Patients

This study was approved by the ethics committees of the above four participant hospitals. Written informed consent was obtained from all participants enrolled in this study. Between March and May in 2020, a total of 54 patients with laboratory confirmed COVID-19 disease were enrolled at the Guangzhou Eighth People’s Hospital, the Shunde Hospital of Guangzhou University of Chinese Medicine, the Fourth People’s Hospital of Foshan, and the First People’s Hospital of Foshan, China. The COVID-19 patients were classified into four groups: 1) The asymptomatic group were defined as patients without self-perception or clinically recognizable symptoms during the whole disease course but diagnosed as COVID-19 according to the Protocol for Prevention and Control of COVID-19 (Edition 6) of National Health and Health Commission of China (37); 2) The symptomatic group were those who had evident clinical symptoms and were diagnosed as COVID-19 according to the Diagnosis and Treatment Protocol for Novel Coronavirus Pneumonia (Version 7) of National Health and Health Commission of China (38); 3) The convalescent group were those who met the discharge standards in the Diagnosis and Treatment Protocol for Novel Coronavirus Pneumonia (Version 7) of National Health and Health Commission of China(38); 4) The re-detectable positive group were defined as patients with re - positive results of SARS-CoV-2 nucleic acid during the follow-up period after discharge from hospital. The 16 health donors were all enrolled at Shunde Hospital of Guangzhou University of Chinese Medicine, China, who tested negative for SARS-CoV-2 and exhibited no respiratory symptoms. Patient demographics and clinical manifestations were retrospectively reviewed. All patients had routine laboratory investigations, including complete blood count, liver function tests, blood gases analysis, and coagulation tests.

### RNA extraction and cDNA synthesis

Peripheral venous blood was collected and placed into the vacutainer tube. The time points of sample collection were shown in Supplementary Table 5. Peripheral blood mononuclear cells (PBMCs) are isolated from 2~4ml human peripheral blood by Ficoll-Paque density gradient. Total RNA was isolated from PBMCs using TRIzol reagent (Invitrogen, USA) according to the manufacturer’s instruction (miRNeasy Mini Kit,Qiagen, Germany).

TCR cDNA libraries for high-throughput sequencing were prepared by 5’rapid amplification of cDNA ends (RACE) using the SMARTScribe™ Reverse Transcriptase (Clontech, USA) as previously described (39, 40). Briefly, 0.6μg of total RNA was mixed with the primer BC1R (Supplementary Table 6), which is specific for human TCRβ cDNA synthesis. To denature RNA and anneal the priming oligonucleotides, RNA was incubated at 70°C for 2 min and then at 42°C for 3 min. Switch_oligo and SMARTScribe reverse transcriptase were added for 25 μl template switching and the cDNA synthesis reaction, which was performed at 42°C for 60 min. 5U uracil DNA glycosylase (UDG) was added for digestion at 37℃ for 40min and product was purified with MinElute PCR Purification Kit (Qiagen, Germany).

### TCR library preparation

Two-round PCR was performed for TCR library preparation. For the first round of PCR amplification, 45 μl of cDNA from the synthesis reaction was mixed with primers and Q5® High-Fidelity 2X Master Mix(NEB, USA). The PCR program began with an initial denaturation at 95°C for 1.5minutes, followed by 18 cycles of denaturation at 95°C for 10s, annealing of primer to DNA at 60°C for 20s, and extension at 72°C for 40s, ended with an extension at 72°C for 4 min. For the second round of PCR amplification, the product from the first round of PCR was purified by QIAquick PCR Purifcation Kit (Qiagen, Germany), 10 μl of the purified product was used in each 25 μl PCR reaction. The reaction was performed for 14 cycles using the first round PCR temperature regimen. PCR products were purified using QIAquick PCR Purifcation Kit (Qiagen, Germany). Illumina adaptors were ligated using NEBNext® Ultra™ II DNA Library Prep Kit for Illumina® (New England BioLabs, USA) according to the manufacturer’s protocol and sequencing in the Illumina platform with the PE150 mode. The parameters of sequencing quality were shown in (Supplementary Table 7).

### Data processing and analysis

The original data obtained from high-throughput sequencing were converted to raw sequence reads by base calling, and the results were stored in FASTQ format. Low-quality reads and reads without primers were discarded. PCR and sequencing errors were corrected by unique molecular identifiers (UMIs). The reads with same UMI were one clone. Only duplicate reads with different UMIs will be kept in downstream processing. A clonotype was defined by the CDR3 amino acid sequence for further analysis and only sequences with a read count ≥2 were included in the analysis. TCRβ V, D, J gene and clonotype defined according to IMGT59 and Igblast; TCRβ VDJ combination defined by MIXCR. We performed mRNA-Seq profiling on 72 samples. Reads mapped to the IPD-IMGT/HLA database and HLA -HD determining HLA alleles using mRNA-seq results (41).

### Statistical analysis

Categorical variables were described as count (%), and two group comparison was performed using Fisher’s exact test. Continuous variables were expressed as median and interquartile range (IQR) values and compared by Mann–Whitney U test between groups. Spearman correlation coefficient was calculated to assess the association between two vectors of quantitative variables. The significance of monotonic trend across multiple groups was assessed by Cuzick’s test (28). The significance of overlap between tow gene sets was assessed by hypergeometric test (the “dhyper” function of R software). All statistical analyses were performed using R software (version 4.0.2). A two-sided P-value lower than 0.05 was considered as statistically significant. Given the exploratory nature of the present study, unadjusted p-values not accounting for multiple testing were used to screen COVID-19 associated TCR variables.

## Acknowledgements

We thank Jinyong He, Yile Huang, Minling Ye from Shunde Hospital of Guangzhou University of Chinese Medicine, Yinong Ye, Weixuan Li from the First People’s Hospital of Foshan, Bingyao Lin, Zhengxing Lin, Jinmei Zhang, Yinzhi Kong from the Forth People’s Hospital of Foshan, and Yanxia Liu, Xing Chen, Mingkai Tan from Guangzhou Eighth People’s Hospital for helping to collect cases and handle samples. We also thank Dan He, Xiaodan Wang, Hongtao Xie and Songbai Zheng from Guangzhou Huayin Medical Laboratory Center for helping with the sequencing experiment.

