## Supplementary tables for "T-cell Repertoire Characteristics of Asymptomatic and Re-detectable Positive COVID-19 Patients"

**Supplementary Table 1.** Blood laboratory tests of study subjects.

|  | **Healthy (n =16)** | **Asymptomatic (n = 11)** | **Symptomatic (n =19 )** | **Convalescent (n =14)** | **Re-detectable positive (n =10)** |
| --- | --- | --- | --- | --- | --- |
| Leukocytes count (×109/L) | 6.71 (6.25-7.49) | 6.18 (5.04-7.23) | 6.49 (5.05-7.14) | 5.27 (4.76-8.18) | 6.24 (4.72-7.80) |
| Neutrophils count (×109/L) | 3.95 (3.56-4.33) | 3.44 (2.73-4.94) | 3.72 (2.54-4.70) | 3.06 (2.36-5.35) | 3.23 (2.79-5.25) |
| Lymphocytes count (×109/L) | 2.25 (1.86-2.41) | 1.66 (1.46-2.24) | 1.72 (1.23-2.12) | 1.70 (1.51-1.96) | 1.89 (1.38-2.39) |
| Monocyte count (×109/L) | 0.41 (0.36-0.48) | 0.36 (0.28-0.60) | 0.42 (0.35-0.49) | 0.37 (0.34-0.70) | 0.29 (0.22-0.33) |
| Eosinophil count (×109/L) | 0.12 (0.09-0.21) | 0.05 (0.03-0.13) | 0.09 (0.04-0.21) | 0.13 (0.08-0.31) | 0.075 (0.055-0.238) |
| Basophils count (×109/L) | 0.03 (0.02-0.05) | 0.03 (0.02-0.04) | 0.01 (0.01-0.03) | 0.015 (0.008-0.023) | 0.02 (0.01-0.03) |
| Erythrocyte count (×109/L) | 5.13 (4.65-5.40) | 4.43 (4.11-4.71) | 4.49 (3.84-5.00) | 4.40 (3.32-4.85) | 4.86 (4.21-5.11) |
| Hemoglobin (g/L） | 144.00 (137.25-159.00) | 134.00 (121.00-148.00) | 131.00 (116.00-149.00) | 131.5 (108.25-149.75) | 142.5 (116-155.5) |
| Platelet count (×109/L) | 263.00 (216.75-295.25) | 258.00 (222.00-290.00) | 256.00 (199.00-288.00) | 244.5 (205.25-283.00) | 243.50 (182.25-301.25) |
| Alanine aminotransferase (ALT,U/L) | 28.55 (24.80-44.38) | 16.00 (13.80-38.40) | 26.30 (18.25-38.60) | 24.73 (13.38-47.85) * | 11.00 (4.75-21.50) |
| Aspartate transaminase (AST,U/L) | 23.75 (20.78-26.03) | 22.80 (17.20-29.00) | 23.00 (13.80-27.90) | 17.65 (13.45-45.00) * | 18.50 (15.00-30.00) |

*Examined in 10 of the convalescent patients.

**Supplementary Table 2.** V-J pairings showing monotonic trend across healthy, asymptomatic and symptomatic groups.

| **V-J Pairing** | **Mean Rank^*^** | | | **P-value** |
| --- | --- | --- | --- | --- |
|  | **Healthy** | **Asymptomatic** | **Symptomatic** |  |
| TRBV11-3 \| TRBJ2-1 | 12.1 | 27.1 | 31.1 | 3.74E-05 |
| TRBV11-1 \| TRBJ2-1 | 14.8 | 20.3 | 32.7 | 7.53E-05 |
| TRBV15 \| TRBJ2-5 | 31.3 | 27.3 | 14.7 | 2.38E-04 |
| TRBV11-1 \| TRBJ2-2 | 15.7 | 21.3 | 31.4 | 5.27E-04 |
| TRBV14 \| TRBJ2-5 | 16.3 | 19.5 | 31.8 | 5.66E-04 |
| TRBV7-6 \| TRBJ2-1 | 13.1 | 29.9 | 28.6 | 8.76E-04 |
| TRBV13 \| TRBJ2-2 | 15.4 | 23.3 | 30.5 | 9.17E-04 |
| TRBV2 \| TRBJ2-3 | 14.3 | 28.5 | 28.3 | 2.60E-03 |
| TRBV5-1 \| TRBJ2-2 | 16.4 | 23.0 | 29.8 | 3.19E-03 |
| TRBV20-1 \| TRBJ1-5 | 31.4 | 21.6 | 17.9 | 3.46E-03 |
| TRBV20/OR9-2 \| TRBJ1-1 | 31.2 | 21.6 | 18.1 | 4.07E-03 |
| TRBV9 \| TRBJ1-2 | 29.6 | 26.3 | 16.7 | 4.23E-03 |
| TRBV11-2 \| TRBJ2-4 | 17.2 | 21.4 | 30.0 | 4.58E-03 |
| TRBV5-1 \| TRBJ2-3 | 15.4 | 26.5 | 28.5 | 4.58E-03 |
| TRBV13 \| TRBJ2-3 | 13.8 | 31.5 | 27.0 | 5.16E-03 |
| TRBV23-1 \| TRBJ2-1 | 14.9 | 28.5 | 27.9 | 5.16E-03 |
| TRBV6-4 \| TRBJ2-2 | 17.4 | 21.1 | 30.0 | 5.36E-03 |
| TRBV4-1 \| TRBJ2-7 | 29.2 | 26.3 | 17.1 | 6.75E-03 |
| TRBV19 \| TRBJ1-5 | 15.6 | 27.3 | 28.0 | 7.29E-03 |
| TRBV20/OR9-2 \| TRBJ2-5 | 28.7 | 27.5 | 16.8 | 7.85E-03 |
| TRBV11-3 \| TRBJ2-2 | 16.1 | 26.0 | 28.3 | 8.15E-03 |
| TRBV5-5 \| TRBJ2-2 | 17.8 | 21.2 | 29.6 | 9.28E-03 |
| TRBV4-2 \| TRBJ2-7 | 29.2 | 25.2 | 17.7 | 1.13E-02 |
| TRBV20-1 \| TRBJ2-7 | 30.5 | 21.3 | 18.9 | 1.18E-02 |
| TRBV7-6 \| TRBJ2-2 | 16.9 | 24.7 | 28.4 | 1.22E-02 |
| TRBV7-9 \| TRBJ2-1 | 18.2 | 21.0 | 29.4 | 1.26E-02 |
| TRBV18 \| TRBJ2-2 | 17.5 | 23.1 | 28.8 | 1.31E-02 |
| TRBV11-2 \| TRBJ2-1 | 16.8 | 25.2 | 28.2 | 1.36E-02 |
| TRBV11-2 \| TRBJ2-2 | 16.2 | 26.9 | 27.6 | 1.40E-02 |
| TRBV29-1 \| TRBJ2-7 | 30.1 | 22.1 | 18.8 | 1.40E-02 |
| TRBV20-1 \| TRBJ2-5 | 29.6 | 23.5 | 18.4 | 1.45E-02 |
| TRBV29-1 \| TRBJ1-5 | 29.2 | 24.5 | 18.1 | 1.48E-02 |
| TRBV25-1 \| TRBJ2-1 | 30.3 | 20.8 | 19.3 | 1.79E-02 |
| TRBV28 \| TRBJ1-4 | 17.6 | 23.5 | 28.4 | 1.79E-02 |
| TRBV25-1 \| TRBJ2-3 | 30.1 | 21.4 | 19.2 | 1.85E-02 |
| TRBV29-1 \| TRBJ2-2 | 19.6 | 17.9 | 30.0 | 1.92E-02 |
| TRBV5-5 \| TRBJ2-5 | 19.4 | 18.7 | 29.7 | 1.98E-02 |
| TRBV6-5 \| TRBJ1-2 | 29.4 | 23.2 | 18.7 | 1.98E-02 |
| TRBV30 \| TRBJ1-5 | 17.9 | 23.0 | 28.5 | 2.05E-02 |
| TRBV14 \| TRBJ2-4 | 16.5 | 27.5 | 27.1 | 2.26E-02 |
| TRBV30 \| TRBJ2-2 | 19.2 | 19.6 | 29.4 | 2.26E-02 |
| TRBV15 \| TRBJ2-7 | 28.9 | 24.0 | 18.7 | 2.50E-02 |
| TRBV15 \| TRBJ2-6 | 30.0 | 20.8 | 19.6 | 2.54E-02 |
| TRBV10-3 \| TRBJ2-6 | 18.8 | 21.1 | 28.8 | 2.58E-02 |
| TRBV5-5 \| TRBJ1-1 | 20.4 | 16.5 | 30.1 | 2.75E-02 |
| TRBV7-9 \| TRBJ1-3 | 18.1 | 23.4 | 28.1 | 2.85E-02 |
| TRBV19 \| TRBJ2-3 | 17.2 | 26.0 | 27.3 | 2.99E-02 |
| TRBV11-1 \| TRBJ2-3 | 17.7 | 24.8 | 27.6 | 3.03E-02 |
| TRBV19 \| TRBJ1-3 | 16.9 | 27.0 | 27.0 | 3.03E-02 |
| TRBV12-3 \| TRBJ2-1 | 30.7 | 18.1 | 20.6 | 3.13E-02 |
| TRBV20/OR9-2 \| TRBJ2-1 | 29.6 | 21.4 | 19.6 | 3.13E-02 |
| TRBV7-8 \| TRBJ2-4 | 17.1 | 26.8 | 27.0 | 3.23E-02 |
| TRBV9 \| TRBJ2-7 | 28.2 | 25.1 | 18.6 | 3.23E-02 |
| TRBV19 \| TRBJ2-2 | 17.7 | 25.0 | 27.5 | 3.33E-02 |
| TRBV5-5 \| TRBJ2-1 | 19.4 | 20.2 | 28.9 | 3.33E-02 |
| TRBV6-1 \| TRBJ2-4 | 29.2 | 22.0 | 19.5 | 3.39E-02 |
| TRBV7-3 \| TRBJ2-3 | 17.2 | 26.7 | 27.0 | 3.44E-02 |
| TRBV7-6 \| TRBJ2-7 | 18.8 | 21.9 | 28.4 | 3.44E-02 |
| TRBV10-3 \| TRBJ2-1 | 18.2 | 23.8 | 27.8 | 3.55E-02 |
| TRBV25-1 \| TRBJ2-5 | 29.8 | 20.1 | 20.2 | 3.78E-02 |
| TRBV27 \| TRBJ2-3 | 29.8 | 20.3 | 20.1 | 3.78E-02 |
| TRBV6-1 \| TRBJ2-7 | 29.8 | 20.3 | 20.1 | 3.78E-02 |
| TRBV5-6 \| TRBJ2-2 | 18.3 | 23.6 | 27.8 | 3.89E-02 |
| TRBV6-6 \| TRBJ1-5 | 20.1 | 18.6 | 29.2 | 3.89E-02 |
| TRBV15 \| TRBJ1-5 | 27.2 | 27.4 | 18.1 | 4.02E-02 |
| TRBV20/OR9-2 \| TRBJ2-7 | 29.6 | 20.6 | 20.1 | 4.02E-02 |
| TRBV2 \| TRBJ2-2 | 20.5 | 17.5 | 29.5 | 4.02E-02 |
| TRBV13 \| TRBJ1-2 | 16.7 | 28.6 | 26.3 | 4.08E-02 |
| TRBV30 \| TRBJ1-3 | 17.2 | 27.4 | 26.6 | 4.33E-02 |
| TRBV5-1 \| TRBJ2-6 | 18.9 | 22.5 | 28.0 | 4.47E-02 |
| TRBV12-5 \| TRBJ2-1 | 21.4 | 15.3 | 30.0 | 4.67E-02 |
| TRBV11-1 \| TRBJ1-1 | 18.5 | 23.9 | 27.5 | 4.88E-02 |
| TRBV12-2 \| TRBJ2-3 | 18.3 | 24.5 | 27.3 | 4.88E-02 |

*Mean rank was calculated as the averaged value of descending rank number.

**Supplementary Table 3.** V-J pairings showing differential usage between healthy and re-detectable positive groups.

| **V-J Pairing** | **Mean Rank^*^** | | **P-value** |
| --- | --- | --- | --- |
|  | **Healthy** | **Re-detectable Positive** |  |
| TRBV13 \| TRBJ1-6 | 17.2 | 7.7 | 1.29E-03 |
| TRBV11-1 \| TRBJ2-3 | 10 | 19.1 | 2.82E-03 |
| TRBV7-8 \| TRBJ1-1 | 10.3 | 18.6 | 4.13E-03 |
| TRBV7-3 \| TRBJ1-1 | 10.7 | 18 | 5.74E-03 |
| TRBV24-1 \| TRBJ2-2 | 10.3 | 18.6 | 6.05E-03 |
| TRBV28 \| TRBJ1-4 | 10.4 | 18.4 | 7.12E-03 |
| TRBV25-1 \| TRBJ2-1 | 16.7 | 8.4 | 7.57E-03 |
| TRBV15 \| TRBJ2-6 | 16.3 | 9.1 | 8.12E-03 |
| TRBV3-1 \| TRBJ1-6 | 16.6 | 8.6 | 9.85E-03 |
| TRBV7-6 \| TRBJ2-1 | 10.5 | 18.3 | 1.03E-02 |
| TRBV11-1 \| TRBJ1-1 | 10.6 | 18.1 | 1.16E-02 |
| TRBV29-1 \| TRBJ1-5 | 16.4 | 8.9 | 1.61E-02 |
| TRBV19 \| TRBJ2-3 | 10.7 | 18 | 1.69E-02 |
| TRBV6-5 \| TRBJ2-6 | 10.7 | 18 | 1.83E-02 |
| TRBV18 \| TRBJ2-2 | 10.7 | 18 | 1.89E-02 |
| TRBV7-6 \| TRBJ2-2 | 10.7 | 18 | 1.90E-02 |
| TRBV19 \| TRBJ1-5 | 10.7 | 18 | 1.96E-02 |
| TRBV11-1 \| TRBJ2-2 | 10.8 | 17.8 | 1.98E-02 |
| TRBV14 \| TRBJ2-4 | 10.8 | 17.8 | 1.98E-02 |
| TRBV29-1 \| TRBJ2-7 | 16.2 | 9.1 | 1.98E-02 |
| TRBV5-1 \| TRBJ2-2 | 10.8 | 17.9 | 1.98E-02 |
| TRBV11-3 \| TRBJ2-5 | 10.8 | 17.8 | 2.11E-02 |
| TRBV7-4 \| TRBJ2-4 | 15 | 11.1 | 2.56E-02 |
| TRBV10-3 \| TRBJ1-3 | 11.2 | 17.2 | 2.60E-02 |
| TRBV15 \| TRBJ2-5 | 16.1 | 9.3 | 2.68E-02 |
| TRBV20-1 \| TRBJ2-7 | 16.1 | 9.3 | 2.68E-02 |
| TRBV7-6 \| TRBJ2-3 | 10.9 | 17.7 | 2.72E-02 |
| TRBV16 \| TRBJ1-3 | 15.6 | 10.2 | 2.83E-02 |
| TRBV11-3 \| TRBJ2-1 | 10.9 | 17.6 | 3.09E-02 |
| TRBV12-3 \| TRBJ2-4 | 11 | 17.5 | 3.56E-02 |
| TRBV18 \| TRBJ1-3 | 11.2 | 17.1 | 3.75E-02 |
| TRBV4-1 \| TRBJ2-4 | 11.1 | 17.4 | 3.83E-02 |
| TRBV6-1 \| TRBJ1-5 | 11.1 | 17.4 | 3.83E-02 |
| TRBV10-3 \| TRBJ2-4 | 11.2 | 17.2 | 4.01E-02 |
| TRBV11-2 \| TRBJ2-3 | 11.1 | 17.4 | 4.08E-02 |
| TRBV5-6 \| TRBJ2-2 | 11.1 | 17.4 | 4.23E-02 |
| TRBV13 \| TRBJ1-2 | 11.1 | 17.3 | 4.60E-02 |
| TRBV11-1 \| TRBJ2-1 | 11.1 | 17.3 | 4.67E-02 |
| TRBV4-2 \| TRBJ2-7 | 15.9 | 9.7 | 4.67E-02 |
| TRBV10-3 \| TRBJ1-1 | 15.9 | 9.7 | 4.80E-02 |

*Mean rank was calculated as the averaged value of descending rank number.

**Supplementary Table 4.** Frequency of HLA alleles.

| **HLA Allele** | **HD Group** | **ASY Group** | **SYM Group** | **CON Group** | **RDP Group** | **P-value^*^** |
| --- | --- | --- | --- | --- | --- | --- |
| **HLA-A** | | | | | | |
| HLA-A*01:01:01 | 2 (6.25%) | 0 (0.00%) | 0 (0.00%) | 0 (0.00%) | 0 (0.00%) | 0.45 |
| HLA-A*02:01:01 | 2 (6.25%) | 4 (18.18%) | 3 (7.89%) | 4 (14.29%) | 5 (25.00%) | 0.45 |
| HLA-A*02:03:01 | 4 (12.50%) | 0 (0.00%) | 0 (0.00%) | 0 (0.00%) | 1 (5.00%) | 0.34 |
| HLA-A*02:05:01 | 0 (0.00%) | 0 (0.00%) | 0 (0.00%) | 0 (0.00%) | 1 (5.00%) | 0.45 |
| HLA-A*02:06:01 | 0 (0.00%) | 1 (4.55%) | 3 (7.89%) | 3 (10.71%) | 0 (0.00%) | 0.45 |
| HLA-A*02:07:01 | 2 (6.25%) | 2 (9.09%) | 1 (2.63%) | 1 (3.57%) | 0 (0.00%) | 0.82 |
| HLA-A*02:09 | 0 (0.00%) | 1 (4.55%) | 0 (0.00%) | 0 (0.00%) | 0 (0.00%) | 0.45 |
| HLA-A*03:01:01 | 0 (0.00%) | 0 (0.00%) | 0 (0.00%) | 0 (0.00%) | 1 (5.00%) | 0.45 |
| HLA-A*11:01:01 | 9 (28.12%) | 4 (18.18%) | 17 (44.74%) | 10 (35.71%) | 7 (35.00%) | 0.45 |
| HLA-A*11:02:01 | 4 (12.50%) | 1 (4.55%) | 1 (2.63%) | 3 (10.71%) | 2 (10.00%) | 0.68 |
| HLA-A*24:02:01 | 4 (12.50%) | 3 (13.64%) | 5 (13.16%) | 2 (7.14%) | 3 (15.00%) | 0.97 |
| HLA-A*24:02:40 | 0 (0.00%) | 0 (0.00%) | 1 (2.63%) | 0 (0.00%) | 0 (0.00%) | 1.00 |
| HLA-A*26:01:01 | 0 (0.00%) | 1 (4.55%) | 0 (0.00%) | 1 (3.57%) | 0 (0.00%) | 0.45 |
| HLA-A*30:01:01 | 0 (0.00%) | 2 (9.09%) | 2 (5.26%) | 3 (10.71%) | 0 (0.00%) | 0.45 |
| HLA-A*30:04:01 | 1 (3.12%) | 0 (0.00%) | 0 (0.00%) | 0 (0.00%) | 0 (0.00%) | 0.82 |
| HLA-A*31:01:02 | 0 (0.00%) | 0 (0.00%) | 3 (7.89%) | 0 (0.00%) | 0 (0.00%) | 0.45 |
| HLA-A*32:01:01 | 0 (0.00%) | 1 (4.55%) | 1 (2.63%) | 0 (0.00%) | 0 (0.00%) | 0.82 |
| HLA-A*33:03:01 | 4 (12.50%) | 2 (9.09%) | 1 (2.63%) | 1 (3.57%) | 0 (0.00%) | 0.45 |
| **HLA-B** | | | | | | |
| HLA-B*07:02:01 | 0 (0.00%) | 1 (4.55%) | 0 (0.00%) | 0 (0.00%) | 0 (0.00%) | 0.72 |
| HLA-B*13:01:01 | 1 (3.12%) | 2 (9.09%) | 0 (0.00%) | 3 (10.71%) | 2 (10.00%) | 0.67 |
| HLA-B*13:02:01 | 1 (3.12%) | 1 (4.55%) | 3 (7.89%) | 3 (10.71%) | 0 (0.00%) | 1.00 |
| HLA-B*14:01:01 | 1 (3.12%) | 0 (0.00%) | 0 (0.00%) | 0 (0.00%) | 0 (0.00%) | 1.00 |
| HLA-B*15:01:01 | 1 (3.12%) | 2 (9.09%) | 1 (2.63%) | 1 (3.57%) | 1 (5.00%) | 1.00 |
| HLA-B*15:02:01 | 6 (18.75%) | 0 (0.00%) | 2 (5.26%) | 2 (7.14%) | 2 (10.00%) | 0.67 |
| HLA-B*15:11:01 | 2 (6.25%) | 0 (0.00%) | 1 (2.63%) | 1 (3.57%) | 2 (10.00%) | 1.00 |
| HLA-B*15:12 | 0 (0.00%) | 1 (4.55%) | 0 (0.00%) | 1 (3.57%) | 0 (0.00%) | 0.72 |
| HLA-B*15:19 | 1 (3.12%) | 0 (0.00%) | 0 (0.00%) | 0 (0.00%) | 0 (0.00%) | 1.00 |
| HLA-B*15:25:01 | 0 (0.00%) | 0 (0.00%) | 0 (0.00%) | 1 (3.57%) | 0 (0.00%) | 1.00 |
| HLA-B*15:27:01 | 0 (0.00%) | 0 (0.00%) | 0 (0.00%) | 0 (0.00%) | 1 (5.00%) | 0.67 |
| HLA-B*18:01:01 | 0 (0.00%) | 0 (0.00%) | 1 (2.63%) | 0 (0.00%) | 0 (0.00%) | 1.00 |
| HLA-B*27:04:01 | 1 (3.12%) | 0 (0.00%) | 2 (5.26%) | 1 (3.57%) | 0 (0.00%) | 1.00 |
| HLA-B*27:24 | 0 (0.00%) | 0 (0.00%) | 1 (2.63%) | 0 (0.00%) | 0 (0.00%) | 1.00 |
| HLA-B*35:01:01 | 0 (0.00%) | 0 (0.00%) | 0 (0.00%) | 1 (3.57%) | 1 (5.00%) | 0.72 |
| HLA-B*35:03:01 | 1 (3.12%) | 1 (4.55%) | 1 (2.63%) | 0 (0.00%) | 0 (0.00%) | 1.00 |
| HLA-B*35:08:01 | 0 (0.00%) | 1 (4.55%) | 0 (0.00%) | 0 (0.00%) | 0 (0.00%) | 0.72 |
| HLA-B*38:02:01 | 2 (6.25%) | 1 (4.55%) | 0 (0.00%) | 1 (3.57%) | 1 (5.00%) | 1.00 |
| HLA-B*39:01:01 | 0 (0.00%) | 0 (0.00%) | 1 (2.63%) | 0 (0.00%) | 0 (0.00%) | 1.00 |
| HLA-B*40:01:02 | 4 (12.50%) | 0 (0.00%) | 12 (31.58%) | 6 (21.43%) | 5 (25.00%) | 0.64 |
| HLA-B*40:02:01 | 1 (3.12%) | 1 (4.55%) | 2 (5.26%) | 0 (0.00%) | 0 (0.00%) | 1.00 |
| HLA-B*40:06:01 | 0 (0.00%) | 0 (0.00%) | 0 (0.00%) | 0 (0.00%) | 1 (5.00%) | 0.67 |
| HLA-B*44:03:01 | 0 (0.00%) | 0 (0.00%) | 0 (0.00%) | 1 (3.57%) | 0 (0.00%) | 1.00 |
| HLA-B*44:03:02 | 0 (0.00%) | 1 (4.55%) | 1 (2.63%) | 0 (0.00%) | 0 (0.00%) | 1.00 |
| HLA-B*46:01:01 | 4 (12.50%) | 2 (9.09%) | 0 (0.00%) | 1 (3.57%) | 0 (0.00%) | 0.67 |
| HLA-B*48:01:01 | 0 (0.00%) | 2 (9.09%) | 2 (5.26%) | 0 (0.00%) | 0 (0.00%) | 0.67 |
| HLA-B*50:01:01 | 0 (0.00%) | 0 (0.00%) | 0 (0.00%) | 0 (0.00%) | 1 (5.00%) | 0.67 |
| HLA-B*51:01:01 | 1 (3.12%) | 1 (4.55%) | 0 (0.00%) | 1 (3.57%) | 0 (0.00%) | 1.00 |
| HLA-B*52:01:01 | 0 (0.00%) | 1 (4.55%) | 3 (7.89%) | 0 (0.00%) | 2 (10.00%) | 0.67 |
| HLA-B*54:01:01 | 0 (0.00%) | 0 (0.00%) | 1 (2.63%) | 0 (0.00%) | 0 (0.00%) | 1.00 |
| HLA-B*55:01:01 | 1 (3.12%) | 0 (0.00%) | 0 (0.00%) | 0 (0.00%) | 0 (0.00%) | 1.00 |
| HLA-B*55:02:01 | 0 (0.00%) | 2 (9.09%) | 1 (2.63%) | 3 (10.71%) | 1 (5.00%) | 0.72 |
| HLA-B*56:01:01 | 1 (3.12%) | 0 (0.00%) | 1 (2.63%) | 0 (0.00%) | 0 (0.00%) | 1.00 |
| HLA-B*57:01:01 | 0 (0.00%) | 0 (0.00%) | 1 (2.63%) | 1 (3.57%) | 0 (0.00%) | 1.00 |
| HLA-B*58:01:01 | 3 (9.38%) | 1 (4.55%) | 1 (2.63%) | 0 (0.00%) | 0 (0.00%) | 0.79 |
| HLA-B*67:01:02 | 0 (0.00%) | 1 (4.55%) | 0 (0.00%) | 0 (0.00%) | 0 (0.00%) | 0.72 |

*FDR adjusted P-value was calculated with Fisher’s exact test.

**Supplementary Table 5.** Time points of sample collection.

| **Sample ID** | **Group** | **Date of admission** | **Date of discharge** | **Date of Re-detectable positive** | **Date of sampling** |
| --- | --- | --- | --- | --- | --- |
| As_1 | ASY | 2020/4/23 | / | / | 2020/4/23 |
| As_2 | ASY | 2020/4/19 | / | / | 2020/4/20 |
| As_3 | ASY | 2020/4/11 | / | / | 2020/4/11 |
| As_4 | ASY | 2020/4/22 | / | / | 2020/4/23 |
| As_5 | ASY | 2020/4/19 | / | / | 2020/4/23 |
| As_6 | ASY | 2020/4/17 | / | / | 2020/4/23 |
| As_7 | ASY | 2020/5/15 | / | / | 2020/5/15 |
| As_8 | ASY | 2020/5/3 | / | / | 2020/5/3 |
| As_9 | ASY | 2020/4/11 | / | / | 2020/4/11 |
| As_10 | ASY | 2020/3/29 | / | / | 2020/4/9 |
| As_11 | ASY | 2020/3/15 | / | / | 2020/4/13 |
| Mi_1 | SYM | 2020/3/29 | / | / | 2020/4/14 |
| Mi_2 | SYM | 2020/3/19 | / | / | 2020/4/11 |
| Mo_1 | SYM | 2020/5/16 | / | / | 2020/5/19 |
| Mo_2 | SYM | 2020/4/18 | / | / | 2020/4/23 |
| Mo_3 | SYM | 2020/4/12 | / | / | 2020/4/23 |
| Mo_4 | SYM | 2020/4/14 | / | / | 2020/4/24 |
| Mo_5 | SYM | 2020/4/8 | / | / | 2020/4/24 |
| Mo_6 | SYM | 2020/6/15 | / | / | 2020/6/15 |
| Mo_7 | SYM | 2020/4/20 | / | / | 2020/4/29 |
| Mo_8 | SYM | 2020/4/26 | / | / | 2020/4/30 |
| Mo_9 | SYM | 2020/4/8 | / | / | 2020/4/30 |
| Mo_10 | SYM | 2020/4/9 | / | / | 2020/4/27 |
| Mo_11 | SYM | 2020/4/20 | / | / | 2020/4/29 |
| Mo_12 | SYM | 2020/3/21 | / | / | 2020/4/10 |
| Mo_13 | SYM | 2020/3/20 | / | / | 2020/4/7 |
| Mo_14 | SYM | 2020/3/29 | / | / | 2020/4/5 |
| Mo_15 | SYM | 2020/3/27 | / | / | 2020/4/10 |
| Mo_16 | SYM | 2020/5/23 | / | / | 2020/5/23 |
| Se-1 | SYM | 2020/5/1 | / | / | 2020/5/8 |
| Re_1 | CON | / | 2020/2/21 | / | 2020/4/23 |
| Re_2 | CON | / | 2020/3/12 | / | 2020/4/23 |
| Re_3 | CON | / | 2020/3/29 | / | 2020/3/29 |
| Re_4 | CON | / | 2020/4/25 | / | 2020/4/25 |
| Re_5 | CON | / | 2020/4/23 | / | 2020/4/23 |
| Re_6 | CON | / | 2020/4/15 | / | 2020/4/15 |
| Re_7 | CON | / | 2020/5/9 | / | 2020/5/8 |
| Re_8 | CON | / | 2020/3/20 | / | 2020/3/25 |
| Re_9 | CON | / | 2020/3/29 | / | 2020/3/29 |
| Re_10 | CON | / | 2020/6/17 | / | 2020/6/16 |
| Re_11 | CON | / | 2020/3/27 | / | 2020/3/27 |
| Re_12 | CON | / | 2020/3/27 | / | 2020/3/27 |
| Re_13 | CON | / | 2020/4/23 | / | 2020/4/30 |
| Re_14 | CON | / | 2020/5/3 | / | 2020/5/3 |
| RP_1 | RDP | / | / | 2020/4/4 | 2020/4/4 |
| RP_2 | RDP | / | / | 2020/3/22 | 2020/4/3 |
| RP_3 | RDP | / | / | 2020/3/29 | 2020/4/4 |
| RP_4 | RDP | / | / | 2020/4/6 | 2020/4/3 |
| RP_5 | RDP | / | / | 2020/3/28 | 2020/4/14 |
| RP_6 | RDP | / | / | 2020/3/29 | 2020/4/13 |
| RP_7 | RDP | / | / | 2020/4/9 | 2020/4/9 |
| RP_8 | RDP | / | / | 2020/4/4 | 2020/4/4 |
| RP_9 | RDP | / | / | 2020/4/16 | 2020/4/16 |
| RP_10 | RDP | / | / | 2020/4/10 | 2020/4/17 |
| Con_1 | HD | / | / | / | 2020/4/22 |
| Con_2 | HD | / | / | / | 2020/4/22 |
| Con_3 | HD | / | / | / | 2020/4/22 |
| Con_4 | HD | / | / | / | 2020/4/22 |
| Con_5 | HD | / | / | / | 2020/4/22 |
| Con_6 | HD | / | / | / | 2020/4/22 |
| Con_7 | HD | / | / | / | 2020/4/22 |
| Con_8 | HD | / | / | / | 2020/4/22 |
| Con_9 | HD | / | / | / | 2020/4/22 |
| Con_10 | HD | / | / | / | 2020/4/23 |
| Con_11 | HD | / | / | / | 2020/4/23 |
| Con_12 | HD | / | / | / | 2020/4/23 |
| Con_13 | HD | / | / | / | 2020/4/23 |
| Con_14 | HD | / | / | / | 2020/4/23 |
| Con_15 | HD | / | / | / | 2020/4/23 |
| Con_16 | HD | / | / | / | 2020/4/23 |

**Supplementary Table 6.** PCR primer sequences.

| **First strand cDNA synthesis** | |
| --- | --- |
| Switch_oligo | AAGCAGUGGTAUCAACGCAGAGUNNNNUNNNNUNNNNUCTT(rG)4 |
| BC1R | CAGTATCTGGAGTCATTGA |
| **First PCR amplification** | |
| Smart20 | AAGCAGTGGTATCAACGCA |
| BC2R | ATTGGGCAGCCCTGATTACACSTTKTTCAGGTCCTC |
| **Second PCR amplification** | |
| Step_1 | (N)4–6(XXXXX)CAGTGGTATCAACGCAGAG |
| Hum bcj | (N)4-6(XXXXX)ATTGGGCAGCCCTGATT |

XXXXX: optional sample barcode. N: Random nucleotides in Unique Molecular Identifiers (UMIs)

**Supplementary Table 7.** Parameters of TCR sequencing quality.

| **Sample** | **Group** | **Raw Reads** | **Clean Reads** | **Reads Utilization (%)** | **Clones** | **Clonotypes** | **Unique V** | **Unique J** | **Unique VJ** |
| --- | --- | --- | --- | --- | --- | --- | --- | --- | --- |
| As_1 | ASY | 27364852 | 11456480 | 83.73 | 2148 | 1978 | 53 | 13 | 363 |
| As_10 | ASY | 17754716 | 8063492 | 90.83 | 2303 | 1628 | 49 | 13 | 323 |
| As_11 | ASY | 27364852 | 12343718 | 90.22 | 5082 | 4434 | 52 | 14 | 465 |
| As_2 | ASY | 15195476 | 6906936 | 90.91 | 2549 | 1583 | 53 | 13 | 342 |
| As_3 | ASY | 27364852 | 12372782 | 90.43 | 4885 | 2679 | 51 | 13 | 404 |
| As_4 | ASY | 27364852 | 12419034 | 90.77 | 3478 | 2858 | 51 | 13 | 421 |
| As_5 | ASY | 27364852 | 11785506 | 86.14 | 1142 | 1001 | 49 | 13 | 319 |
| As_6 | ASY | 27364852 | 11958368 | 87.4 | 7798 | 2456 | 51 | 13 | 351 |
| As_7 | ASY | 21219360 | 9683912 | 91.27 | 2422 | 1712 | 46 | 13 | 347 |
| As_8 | ASY | 27364852 | 11245632 | 82.19 | 1617 | 1425 | 48 | 13 | 320 |
| As_9 | ASY | 27364852 | 12037324 | 87.98 | 2810 | 1605 | 51 | 14 | 345 |
| Con_1 | HD | 27364852 | 11184432 | 81.74 | 2661 | 1608 | 51 | 13 | 338 |
| Con_10 | HD | 27364852 | 12433020 | 90.87 | 4255 | 3290 | 55 | 14 | 449 |
| Con_11 | HD | 27364852 | 12353914 | 90.29 | 5673 | 4685 | 53 | 14 | 457 |
| Con_12 | HD | 21411044 | 8903966 | 83.17 | 2168 | 1591 | 56 | 14 | 357 |
| Con_13 | HD | 19516052 | 8148356 | 83.5 | 2893 | 1376 | 51 | 14 | 334 |
| Con_14 | HD | 27364852 | 12429334 | 90.84 | 4485 | 3205 | 53 | 13 | 447 |
| Con_15 | HD | 27364852 | 12373566 | 90.43 | 5161 | 3042 | 51 | 13 | 416 |
| Con_16 | HD | 27364852 | 12372016 | 90.42 | 5002 | 3460 | 51 | 13 | 439 |
| Con_2 | HD | 18402992 | 7636542 | 82.99 | 2812 | 1957 | 49 | 13 | 355 |
| Con_3 | HD | 23238312 | 9736980 | 83.8 | 2598 | 2162 | 50 | 14 | 387 |
| Con_4 | HD | 26018584 | 11367410 | 87.38 | 1772 | 1070 | 48 | 13 | 303 |
| Con_5 | HD | 27364852 | 11766710 | 86 | 1667 | 1422 | 51 | 13 | 329 |
| Con_6 | HD | 25640692 | 10636490 | 82.97 | 2408 | 1348 | 49 | 13 | 336 |
| Con_7 | HD | 16982100 | 7081708 | 83.4 | 2833 | 1152 | 50 | 13 | 340 |
| Con_8 | HD | 25547936 | 10963950 | 85.83 | 2447 | 1332 | 51 | 13 | 339 |
| Con_9 | HD | 19809792 | 8186708 | 82.65 | 1468 | 1250 | 51 | 13 | 324 |
| Mi_1 | SYM | 27364852 | 12452246 | 91.01 | 4435 | 3575 | 54 | 13 | 438 |
| Mi_2 | SYM | 22006684 | 10031544 | 91.17 | 2103 | 1858 | 53 | 13 | 344 |
| Mo_1 | SYM | 21516812 | 9816316 | 91.24 | 2504 | 1900 | 49 | 13 | 346 |
| Mo_10 | SYM | 27364852 | 11805250 | 86.28 | 2791 | 1699 | 49 | 13 | 326 |
| Mo_11 | SYM | 27364852 | 11891688 | 86.91 | 2971 | 1600 | 49 | 13 | 319 |
| Mo_12 | SYM | 18188780 | 8246124 | 90.67 | 2553 | 1481 | 51 | 13 | 345 |
| Mo_13 | SYM | 28155936 | 12822126 | 91.08 | 2727 | 1431 | 48 | 13 | 308 |
| Mo_14 | SYM | 27364852 | 11287288 | 82.49 | 3184 | 2621 | 50 | 14 | 396 |
| Mo_15 | SYM | 27364852 | 11267000 | 82.35 | 3281 | 2772 | 49 | 14 | 401 |
| Mo_16 | SYM | 19787344 | 9049988 | 91.47 | 2229 | 1705 | 49 | 14 | 348 |
| Mo_2 | SYM | 14864492 | 6790932 | 91.37 | 2870 | 1611 | 49 | 14 | 336 |
| Mo_3 | SYM | 19729488 | 8953844 | 90.77 | 2089 | 1685 | 48 | 13 | 329 |
| Mo_4 | SYM | 27364852 | 11790628 | 86.17 | 1373 | 1019 | 45 | 13 | 290 |
| Mo_5 | SYM | 27364852 | 11834328 | 86.49 | 1586 | 1453 | 46 | 13 | 314 |
| Mo_6 | SYM | 16795228 | 7581858 | 90.29 | 2926 | 1571 | 50 | 13 | 317 |
| Mo_7 | SYM | 27364852 | 11849956 | 86.61 | 2130 | 1554 | 49 | 13 | 319 |
| Mo_8 | SYM | 27364852 | 11833170 | 86.48 | 2434 | 1917 | 53 | 13 | 338 |
| Mo_9 | SYM | 27364852 | 11900546 | 86.98 | 2924 | 2389 | 49 | 13 | 402 |
| Re_1 | CON | 27364852 | 12143284 | 88.75 | 7267 | 4241 | 51 | 14 | 442 |
| Re_10 | CON | 17826748 | 8141870 | 91.34 | 3205 | 2468 | 53 | 13 | 389 |
| Re_11 | CON | 22181920 | 7856620 | 70.84 | 1873 | 1678 | 51 | 13 | 355 |
| Re_12 | CON | 22845464 | 8987344 | 78.68 | 3039 | 2364 | 50 | 13 | 435 |
| Re_13 | CON | 28852840 | 13019172 | 90.25 | 2442 | 1936 | 51 | 13 | 336 |
| Re_14 | CON | 21590564 | 9806508 | 90.84 | 1928 | 1542 | 49 | 13 | 337 |
| Re_2 | CON | 27364852 | 12377756 | 90.46 | 3309 | 931 | 46 | 14 | 268 |
| Re_3 | CON | 26302468 | 10679048 | 81.2 | 2223 | 1749 | 52 | 13 | 368 |
| Re_4 | CON | 18512268 | 7242544 | 78.25 | 3828 | 2330 | 51 | 13 | 421 |
| Re_5 | CON | 22522024 | 8781726 | 77.98 | 3229 | 2831 | 55 | 13 | 435 |
| Re_6 | CON | 23831776 | 9689068 | 81.31 | 2823 | 2116 | 50 | 13 | 416 |
| Re_7 | CON | 28853344 | 13088104 | 90.72 | 2011 | 1492 | 50 | 14 | 305 |
| Re_8 | CON | 23275636 | 10577948 | 90.89 | 2832 | 712 | 45 | 13 | 241 |
| Re_9 | CON | 22600720 | 9231298 | 81.69 | 2781 | 1908 | 52 | 13 | 377 |
| RP_1 | RDP | 28821976 | 13114332 | 91 | 2937 | 1704 | 52 | 13 | 335 |
| RP_10 | RDP | 27364852 | 12480046 | 91.21 | 4121 | 3109 | 51 | 13 | 420 |
| RP_2 | RDP | 28803420 | 13123492 | 91.12 | 2142 | 1844 | 50 | 13 | 334 |
| RP_3 | RDP | 28855836 | 13179912 | 91.35 | 1908 | 1172 | 47 | 14 | 267 |
| RP_4 | RDP | 28835568 | 13182924 | 91.44 | 2298 | 1623 | 46 | 13 | 351 |
| RP_5 | RDP | 17388316 | 7925718 | 91.16 | 3166 | 1645 | 49 | 13 | 338 |
| RP_6 | RDP | 26477152 | 11944476 | 90.22 | 4652 | 3450 | 50 | 13 | 404 |
| RP_7 | RDP | 13637324 | 6179930 | 90.63 | 1724 | 1336 | 48 | 13 | 295 |
| RP_8 | RDP | 22452948 | 10239876 | 91.21 | 2227 | 1509 | 54 | 13 | 325 |
| RP_9 | RDP | 27364852 | 12356872 | 90.31 | 5024 | 2571 | 52 | 13 | 401 |
| Se_1 | SYM | 18018396 | 8199810 | 91.02 | 4265 | 1528 | 46 | 14 | 331 |
